# Exosomes derived from BMSCs with miR-124-3p inhibitor protects against LPS-induced endometritis through regulation of DUSP6, p-p65 and p-ERK

**DOI:** 10.1101/2023.02.02.526752

**Authors:** Yihong Chen, Shan Zheng, Xiumei Zhao, Yi Zhang, Suchai Yu, Juanbing Wei

## Abstract

Endometritis seriously affects women’s normal life and work. It has been found that microRNA-123-3p (miR-124-3p) expression is abnormally high expression in the patients of chronic endometritis. However, the underlying mechanism for miR-124-3p regulation of endometritis development remains unclear. In our study, we treated human endometrial epithelial cells (HEECs) with LPS to simulate endometrial injury *in vitro*. Then, HEEC was treated with miR-124-3p mimics and miR-124-3p inhibitor. Next, exosomes were separated from bone marrow-derived mesenchymal stem cells (BMSCs). In addition, BMSCs were co-cultured with HEEC. Later on, dual-luciferase reporter assay was carried out to validate the regulation between miR-124-3p and DUSP6. Results indicated that LPS inhibited the viability of HEEC in time and dose dependent manner. MiR-124-3p inhibitor reversed apoptosis and viability inhibition of HEEC which were induced by LPS. In addition, we also found exosomes could transfer miR-124-3p from BMSCs to HEEC. Besides, BMSCs/anti-miR-124-3p Exo was observed to abolish LPS-induced viability and proliferation inhibition of HEEC by inducing the apoptosis of HEEC. Moreover, BMSCs/anti-miR-124-3p Exo alleviated inflammation of HEEC induced by LPS via upregulating DUSP6 and downregulating p-p65 and p-ERK. Furthermore, BMSCs/anti-miR-124-3p Exo protected against LPS-induced endometritis *in vivo* by upregulating DUSP6 and downregulating p-p65 and p-ERK. In conclusion, we found that BMSCs/anti-miR-124-3p Exo might be a promising new alternative to treat endometritis.

## Introduction

Endometritis is one of common gynecological diseases in women of childbearing age and is an inflammatory change in the endometrial structure caused by various reasons ^1–3^. The nosogenesis of endometritis include the endocrine changes, the invasion of exogenous pathogens (tuberculosis, pathogenic bacteria and so on) and age^4–6^. The manifestations in patients with endometritis include lower abdominal pain, mild fever, and increased vaginal discharge^7,8^. Endometritis is often divided into acute endometritis and chronic endometritis^9,10^. Chronic endometritis is the most common cause of miscarriage^11,12^. At present, the main treatment methods for endometritis are antibiotic treatment, traditional Chinese medicine treatment, drug treatment and surgical treatment^13,14^. Nevertheless, the effects of these therapies remain unsatisfactory.

MicroRNAs (miRNAs) are non-coding RNAs with 20-25 nucleotides in length and play regulatory functions in eukaryotes^15,16^. MiRNAs can degrade target mRNA or repress translation of target mRNA^17^. MiRNAs is extensively involved in development endometritis^18,19^. For example, the upregulation of miR-148a can alleviate inflammation of endometrial epithelial cells which are induced by LPS^19^. Additionally, miR-193a-3p overexpression promotes inflammation of bovine endometrial epithelial cells (bEECs) induced by LPS^20^. Moreover, miR-124-3p expression level is abnormally high in the patients of chronic endometritis, compared with the healthy women^18^. However, the mechanism underlying miR-124-3p regulation of endometritis development remains unclear.

BMSCs are a type of adult stem cells with their capacity for high self-renewal and multi-directional differentiation^21^. Under certain conditions, BMSCs can differentiate into a variety of tissue cells, and has the functions of hematopoietic support, immune regulation, tissue repair and so on^22,23^. Exosomes are discoid vesicles with diameters of 40-150 nm. Exosomes derived from BMSCs are widely used in tissue repair^24^. Nevertheless, whether miR-124-3p could be transferred from BMSCs to Human endometrial epithelial cell (HEEC) by exosomes remains unclear.

## Material and methods

### Cell culture

HEEC was provided by Procell Life Science &Technology Co., Ltd. Bone BMSCs were commercially available from ATCC. DMEM contained with 10% fetal bovine serum (FBS), 1% penicillin-streptomycin was purchase from Thermo Fisher. HEEC and BMSCs were cultured in DMEM at 37°C, 5% CO_2_.

### Cell Counting Kit-8 (CCK-8) assay

CCK-8 was used to assess the viability of HEEC at 0, 6, 12 and 24 h. Firstly, HEEC was treated with 0, 0.1, 0.5, 1 or 2 μg/mL LPS, followed by incubation with CCK-8 reagent for 2 h. Later on, the absorbance at 450 nm was measured by a microplate reader. CCK-8 reagent was provided by Beyotime.

### Cell transfection

MiR-124-3p negative control (NC), miR-124-3p mimics, miR-124-3p inhibitor, DUSP6 siRNA1, DUSP6 siRNA2 and DUSP6 siRNA control (siRNA-ctrl) were provided by Genepharma. NC, miR-124-3p mimics, miR-124-3p inhibitor, DUSP6 siRNA1, DUSP6 siRNA2 and DUSP6 siRNA control (siRNA-ctrl) were transfected into HEEC or BMSCs using Lipofectamine® 2000, following the manufacturer’s protocol.

### Reverse transcription-quantitative polymerase chain reaction (RT-qPCR)

Trizol reagent was used to extract total RNA of HEEC, BMSCs or tissue. To obtain cDNA, EntiLink™ 1st Strand cDNA Synthesis Kit was performed. Then, EnTurbo™ SYBR Green PCR SuperMix was performed for qPCR. All reagents were provided by ELK Biotechnology. The data of RT-qPCR was determined based on the 2^−ΔΔCt^ method.

### Flow cytometry

We used flow cytometry to analyze apoptosis of HEEC. Firstly, HEEC was treated with 2 μg/mL LPS, miR-124-3p inhibitor NC and miR-124-3p inhibitor. Then, HEEC were cultured in 6-well plate. After that, HEEC were incubated with Annexin V-FITC and followed by PI for another 15 min. Later on, the apoptosis of HEEC was tested by flow cytometry.

### Enzyme Linked Immune Sorbent Assay (ELISA)

Human IL-10 ELISA kit, human IL6 (Interleukin 6) ELISA kit and Human TGF-β ELISA kit were provided by ELK Biotechnology. Firstly, the supernatant of HEEC and tissue were collected after treatment. Then, the expressions of IL-6, IL-1β and TNF-α in the supernatant of HEEC and tissue were analyzed through according kits respectively, following the manufacturer’s procedure,

### Exosome labeling and uptake

Firstly, exosomes derived from BMSCs that was treated with miR-124-3p (BMSCs/NC Exo) and exosomes derived from BMSCs that was treated with miR-124-3p inhibitor (BMSCs/anti-miR-124-3p Exo) using ultracentrifugation method. The isolated vesicles were identified using Transmission electron microscopy (TEM) and Nanoparticle Tracking Analysis (NTA). In addition, the expressions of TSG101 and CD81 in the isolated vesicles were assessed using western blot. Then, the collected exosomes were labeled using PKH26. Cytoskeleton was labeled using phalloidin. Cell nucleus was labeled using DAPI. Later on, BMSCs were co-cultured with HEEC. Fluorescence microscope was performed to observe whether exosomes could be transferred from BMSCs to HEEC.

### Western blotting

We used RIPA buffer to isolate total protein. RIPA buffer was provided by Aspen Biotechnology. After that, BCA protein assay kit was performed for protein quantification. Then, protein was separated with 10% sodium dodecyl sulfate-polyacrylamide gel electrophoresis (SDS-PAGE), transferred to PVDF membrane and followed by primary antibodies incubation overnight at 4 °C. Next, protein was incubated with HRP-conjugated secondary antibodies for 1 h at 37°C. Finally, ECL kit was used to observe protein bands. Primary antibodies (anti-p-p65, anti-TSG101, anti-CD81, anti-DUSP6, anti-p-ERK1/2, anti-p65 and anti-ERK1/2) and secondary antibodies were provided by Abcam.

### 5-ethynyl-2’-deoxyuridine (EdU) staining

Firstly, HEEC was treated with LPS, LPS + BMSCs/NC Exo or LPS + BMSCs/anti-miR-124-3p Exo. After incubation with 50 μM EdU for 1 h, HEECs were incubated with 1 mg/mL DAPI for 10 min at 37 °C. Later that, the staining results was observed using a microscope. EdU Detection kit was provided by Ribobio.

### Dual-luciferase reporter assay

HEEC were co-transfected with pGL6-miRNA luciferase reporter vector containing DUSP6 wild type (WT) or mutant type (MT), together with miR-124-3p mimics or NC using Lipofectamine 2000. Then, the association between miR-124-3p and DUSP6 was determined by dual-luciferase reporter assay. The pGL6-miRNA luciferase reporter vector was provided by Beyotime.

### Animal study

BALB/c female virgin mice were 8-week-old and provided by Vital River. We randomly separated mice into 4 groups: control, LPS, LPS + BMSCs/NC Exo and LPS + BMSCs/anti-miR-124-3p Exo. The mice were injected with BMSCs/NC Exo and BMSCs/anti-miR-124-3p Exo in utero for 3 days. Three days later, to establish of endometrial injury, the mice were infused with 1 mg/kg LPS for 24 h on each side of the uterus^25,26^. The control group mice were treated with PBS for 1 day. Then, mice were euthanasia and tissue were collected at the end of study. HE and Masson staining were performed to observe LPS induced endometritis in tissue. This study was approved by the National Institutes of Health Guide for the Care and Use of laboratory animals.

### Statistical analysis

Statistical data was analyzed by GraphPad Prism. In each figure legends, the statistical results were shown as mean ± SD. To compare the difference between three or more group, one-way analysis of variance (ANOVA) and Tukey’s test. If P < 0.05, the results were defined as statistically significant.

## Results

### MiR-124-3p inhibitor reverses LPS-induced apoptosis and viability inhibition of HEEC

To study the effect of different dose of LPS on the viability of HEEC, CCK-8 was performed. We found that LPS inhibited the viability of HEEC in time and dose dependent manner (Fig. 1A). Because when the concentration of LPS was 2 μg/mL, the viability of HEEC was about 50%. Therefore, 2 μg/mL LPS was used for the following study. LPS increased miR-124-3p expression in HEEC (Fig. 1B). At the same time, miR-124-3p mimics upregulated miR-124-3p expression in HEEC (Fig. 1C). Expectedly, miR-124-3p inhibitor downregulated the miR-124-3p expression in HEEC (Fig. 1C). Moreover, LPS inhibited the viability of HEEC by inducing the apoptosis, and the effect of LPS was obligated by miR-124-3p inhibitor (Fig. 1D and 1E). Furthermore, ELISA results were shown that LPS promoted the expressions of IL-6 and IL-1β in HEEC, while these phenomena were reversed by miR-124-3p inhibitor (Fig. 1F and 1G). Collectively, inhibition of miR-124-3p reversed LPS-induced apoptosis and viability inhibition in HEEC.

**Figure 1.**
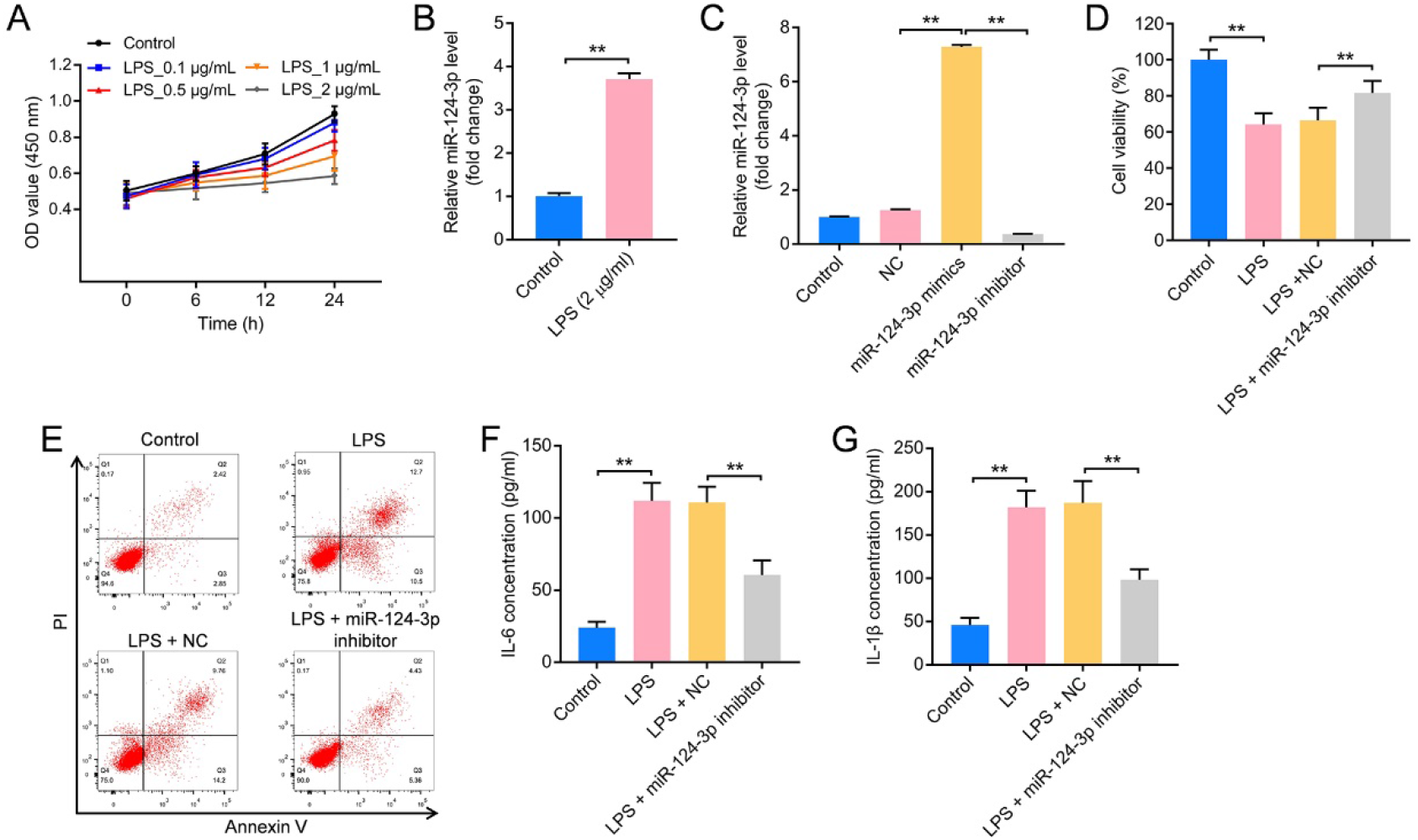
MiR-124-3p inhibitor reverses LPS-induced apoptosis and viability inhibition of HEEC. **(A)** HEEC was treated with 0, 0.1, 0.5, 1 and 2 μg/mL LPS. The viability of HEEC at 0, 6, 12 and 24 h was assessed using CCK-8. **(B)** The expression of miR-124-3p in HEEC was assessed using RT-qPCR. **(C)** HEEC was treated with miR-124-3p mimics and miR-124-3p inhibitor. The expression of miR-124-3p in HEEC was assessed using RT-qPCR. **(D)** The viability of HEEC was assessed using CCK-8. **(E)** The apoptosis of HEEC was assessed using flow cytometry. **(F and G)** The expressions of IL-6 and IL-1β in HEEC were assessed using ELISA. **P < 0.01. n = 3.

### MiR-124-3p is transferred from BMSCs to HEEC by exosomes

It has been reported that BMSCs have the function of repairing tissue damage by delivering exosomes. Therefore, we next explored whether exosome can transfer miR-124-3p from BMSCs to HEEC. Firstly, exosomes were isolated from BMSCs. As shown in Fig. 2A, the diameter of isolated vesicles was 40-150 nm. Besides, the isolated vesicles had a phospholipid bilayer structure (Fig. 2B). Additionally, the isolated vesicles could express the specific markers for exosomes (TSG101 and CD81) (Fig. 2C). These phenomena implied that the isolated vesicles were exosomes. Next, we studied whether exosome can transfer miR-124-3p from BMSCs to HEEC. Firstly, miR-124-3p inhibitor inhibited the miR-124-3p expression level in BMSCs (Fig. 2D). Meanwhile, miR-124-3p was downregulated in exosomes derived from BMSCs that was treated with miR-124-3p (BMSCs/NC Exo) than it was in exosomes derived from BMSCs that was treated with miR-124-3p inhibitor (BMSCs/anti-miR-124-3p Exo) (Fig. 2E). Additionally, PKH26-labeled exosomes could be observed in HEEC (Fig. 2F). These phenomena indicated that exosomes could be transferred from BMSCs to HEEC.

**Figure 2.**
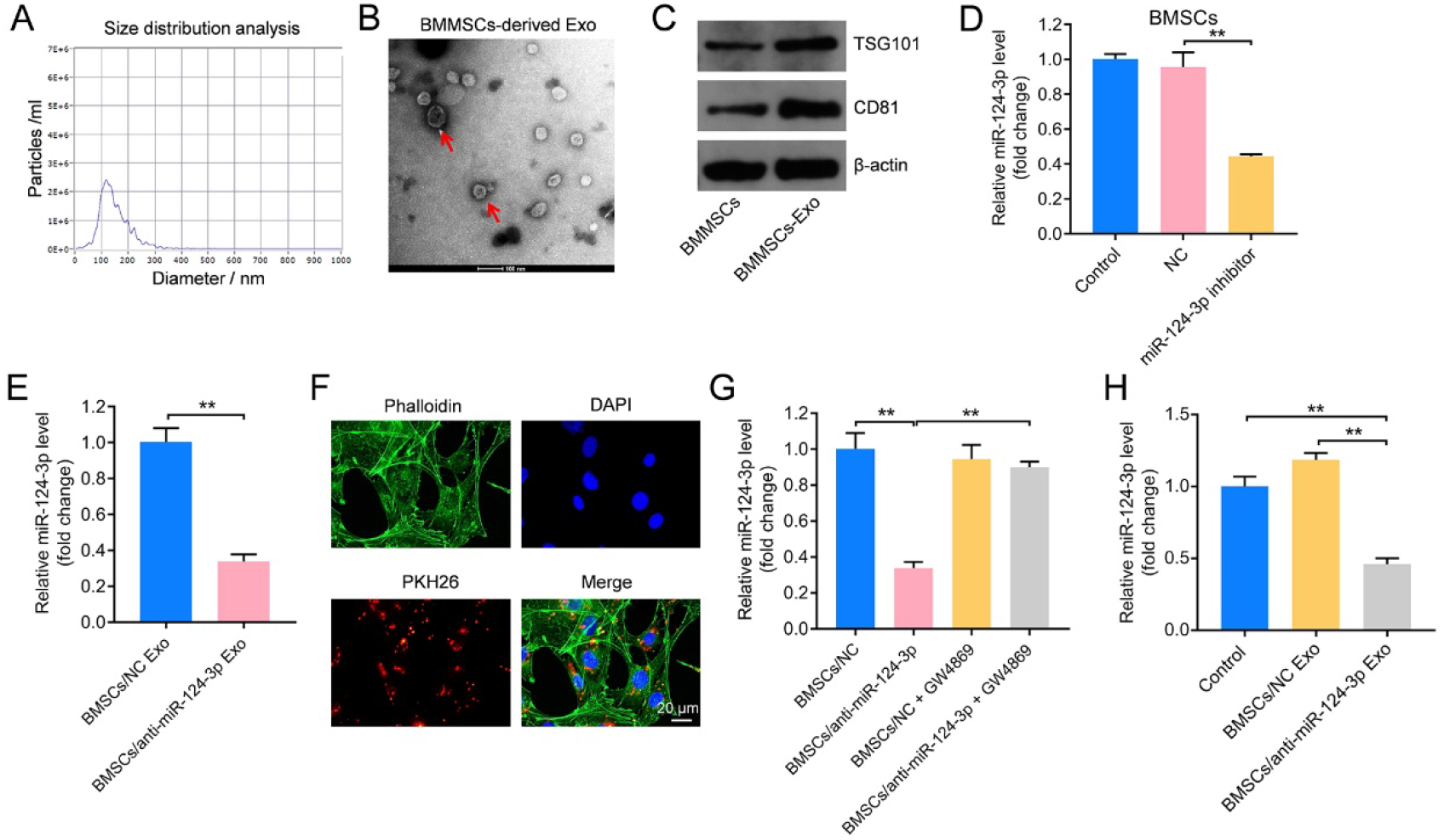
MiR-124-3p is transferred from BMSCs to HEEC by exosomes. **(A and B)** The isolated vesicles were identified using NTA and TEM. **(C)** The expressions of TSG101 and CD81 were assessed using western blot. **(D)** The expression of miR-124-3p in BMSCs was assessed using RT-qPCR. **(E)** The expression of miR-124-3p in exosomes was assessed using RT-qPCR. **(F)** Exosomes were labeled using PKH26. BMSCs were co-cultured with HEEC. Fluorescence microscope was performed to observe whether exosomes could be transferred from BMSCs to HEEC. **(G and H)** The expression of miR-124-3p in HEEC was assessed using RT-qPCR. **P < 0.01. n = 3.

Next, the results of RT-qPCR indicated that BMSCs/anti-miR-124-3p downregulated miR-124-3p expression level in HEEC that was co-cultured with BMSCs. This decrease was reserved by the inhibitor of exosomes, GW4869. (Fig. 2G). In addition, BMSCs/anti-miR-124-3p Exo downregulated miR-124-3p expression level in HEEC (Fig. 2H). Collectively, exosomes can transfer miR-124-3p from BMSCs to HEEC.

### BMSCs/anti-miR-124-3p Exo abolishes LPS-induced viability and proliferation inhibition of HEEC by inducing the apoptosis of HEEC

To explore the role of BMSCs/anti-miR-124-3p Exo on the viability of HEEC, CCK-8 was performed. We found that LPS inhibited the viability and proliferation of HEEC (Fig. 3A and 3B). However, these inhibitions were eliminated by BMSCs/anti-miR-124-3p Exo. Meanwhile, LPS increased the levels of Cleaved caspase 3 and Bax in HEEC, and decreased the level of Bcl-2. These phenomena were reserved by BMSCs/anti-miR-124-3p Exo (Fig. 3C, 3D and 3E). Collectively, BMSCs/anti-miR-124-3p Exo abolished LPS-induced viability and proliferation inhibition of HEEC by inducing the apoptosis of HEEC.

**Figure 3.**
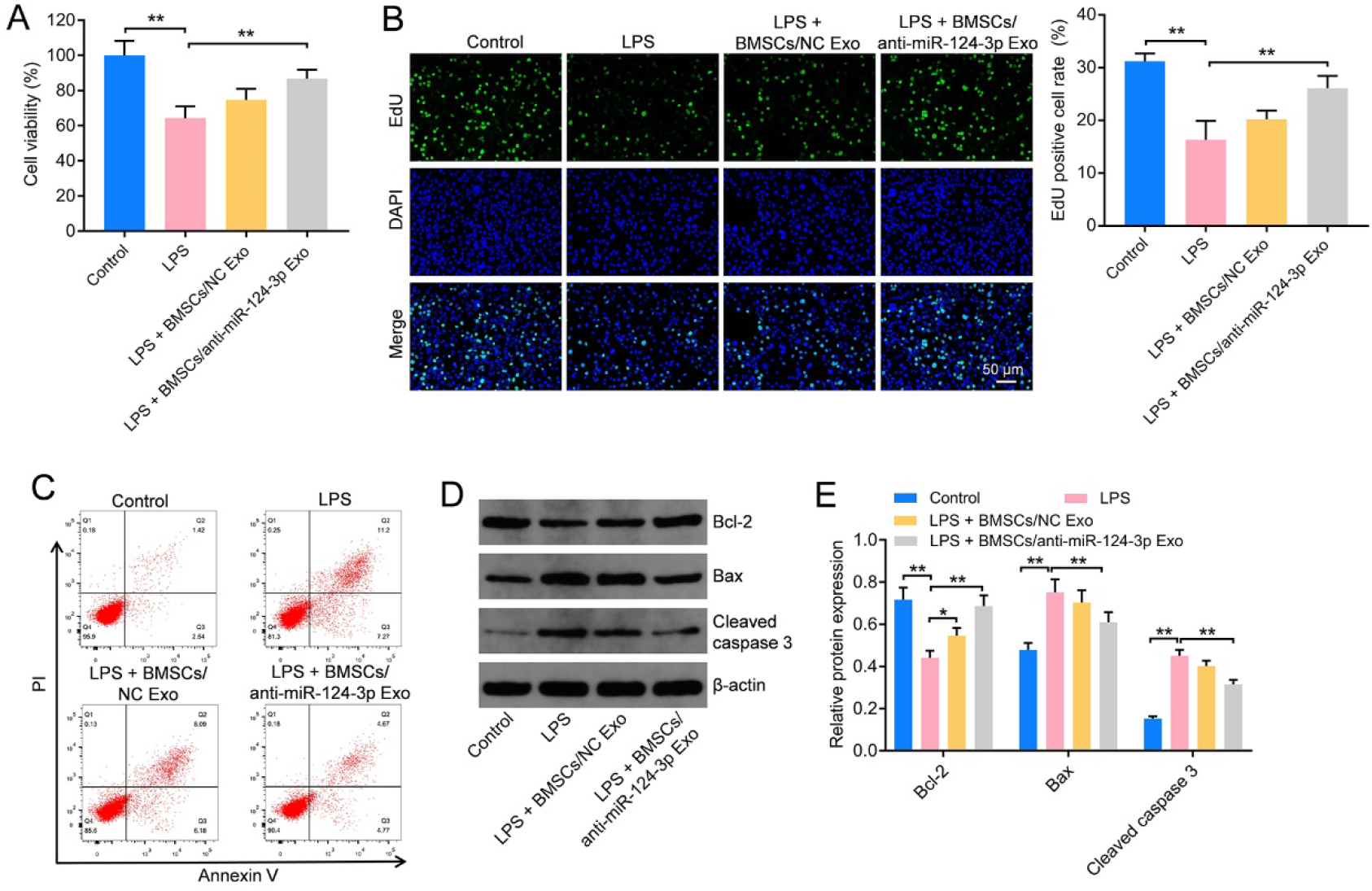
BMSCs/anti-miR-124-3p Exo abolishes LPS-induced viability and proliferation inhibition of HEEC by inducing the apoptosis of HEEC. **(A)** The viability of HEEC was assessed using CCK-8. **(B)** The proliferation of HEEC was assessed using EdU staining. **(C)** The apoptosis of HEEC was assessed using flow cytometry. **(D and E)** The levels of Bcl-2, Bax and Cleaved caspase 3 in HEEC were assessed using western blot. **P < 0.01. n = 3.

### DUSP6 is targeting miR-124-3p

ELISA was performed to assess the role of BMSCs/anti-miR-124-3p Exo on the inflammation of HEEC. As illustrated in Fig. 4A, LPS promoted the expressions of IL-6, TNF-α and IL-1β in HEEC, and these promotions were reserved by BMSCs/anti-miR-124-3p Exo. Next, in order to investigate the mechanism for miR-124-3p regulation of endometritis development, Targetscan database was used. The online database showed that DUSP6 might be a potential target of miR-124-3p (Fig. 4B). Additionally, it has been reported that DUSP6 and endometrium are closely related. Thus, DUSP6 was used for the following study. Dual-luciferase reporter assay validated that miR-124-3p mimics downregulated the luciferase activity of HEEC harboring of DUSP6 WT (Fig. 4C) but not the luciferase activity of HEEC harboring mutant type of DUSP6 (Fig. 4C). Meanwhile, miR-124-3p inhibitor enhanced DUSP6 level (Fig. 4D). Expectedly, miR-124-3p mimics decreased DUSP6 level in HEEC, and miR-124-3p inhibitor upregulated DUSP6 level (Fig. 4E). Collectively, BMSCs/anti-miR-124-3p Exo alleviated inflammation of HEEC induced by LPS. In addition, DUSP6 was targeting miR-124-3p.

**Figure 4.**
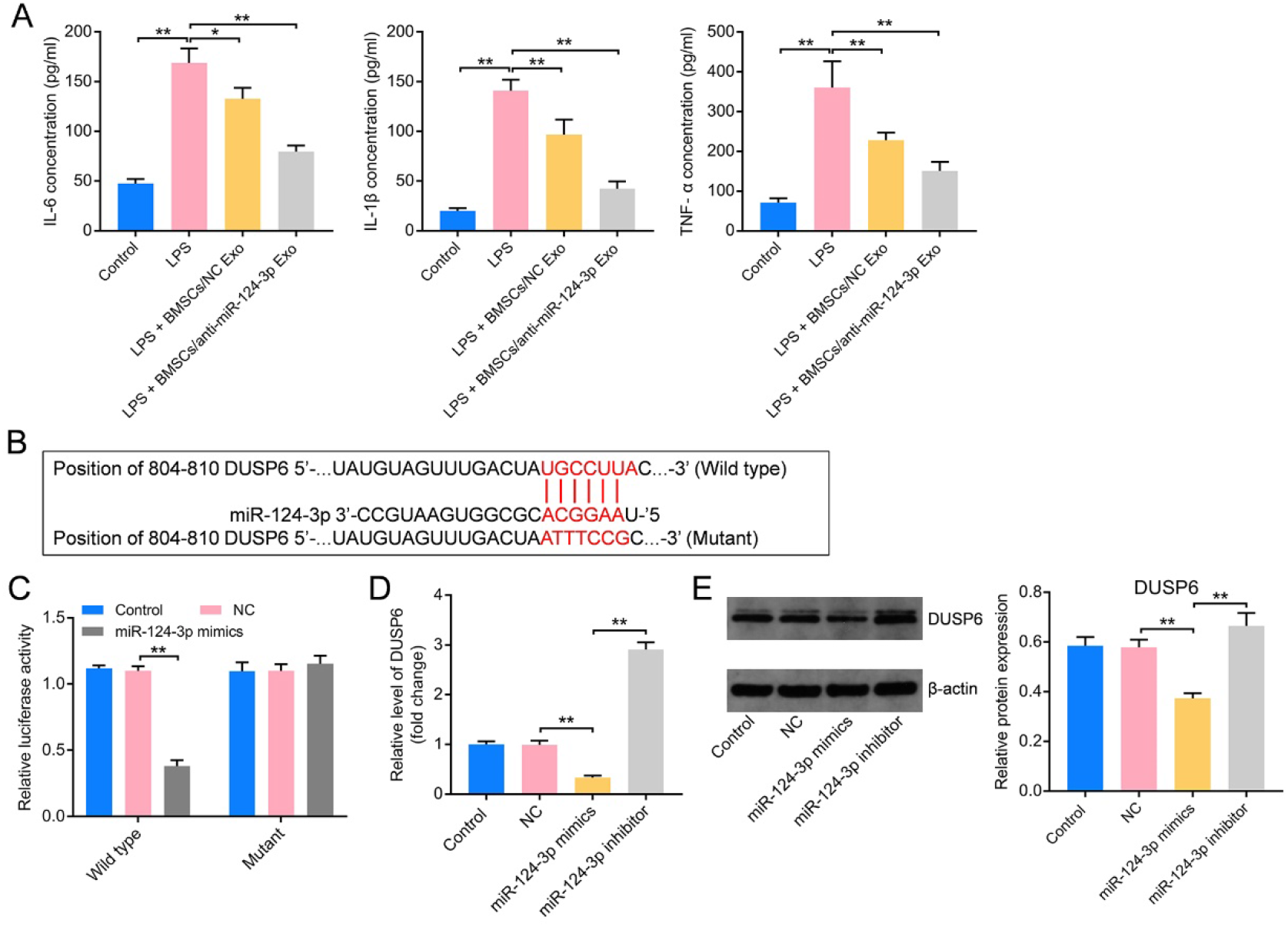
DUSP6 was a target of miR-124-3p. **(A)** The expressions of IL-6, IL-1β and TNF-α in HEEC were assessed using ELISA. **(B)** Targetscan database was used to predict DUSP6 might be one of the targets of miR-124-3p. **(C)** Dual-luciferase reporter assay was performed to confirm the relationship between miR-124-3p and DUSP6. **(D)** The expression of DUSP6 in HEEC was assessed using RT-qPCR. **(E)** The expression of DUSP6 in HEEC was assessed using western blot. **P < 0.01. n = 3.

### BMSCs/anti-miR-124-3p Exo alleviates LPS-induced inflammation of HEEC by upregulating DUSP6 and downregulating p-p65 and p-ERK

To study the role of DUSP6 on HEEC, DUSP6 was knockdown. As indicated in Fig. 5A, DUSP6 siRNAs decreased the level of DUSP6 in HEEC. Because DUSP6 siRNA2 showed the better knockdown effect, DUSP6 siRNA2 was used for following study (Fig. 5A). In addition, LPS inhibited the viability of HEEC, and this inhibition was eliminated by BMSCs/anti-miR-124-3p Exo. However, BMSCs/anti-miR-124-3p Exo effect was reserved by DUSP6 siRNA2 (Fig. 5B). Besides, LPS promoted the expressions of IL-6, TNF-α and IL-1β in HEEC, and these promotions were reserved by BMSCs/anti-miR-124-3p Exo, while, the effect of BMSCs/anti-miR-124-3p Exo was abolished by DUSP6 (Fig. 5C). Moreover, LPS downregulated DUSP6 level in HEEC, and upregulated p-p65 and p-ERK levels; while these phenomena were reserved by BMSCs/anti-miR-124-3p Exo. Nevertheless, the effect of BMSCs/anti-miR-124-3p Exo on these proteins was eliminated by DUSP6 (Fig. 5D). Collectively, BMSCs/anti-miR-124-3p Exo alleviated LPS-induced inflammation of HEEC by upregulating DUSP6 and downregulating p-p65 and p-ERK.

**Figure 5.**
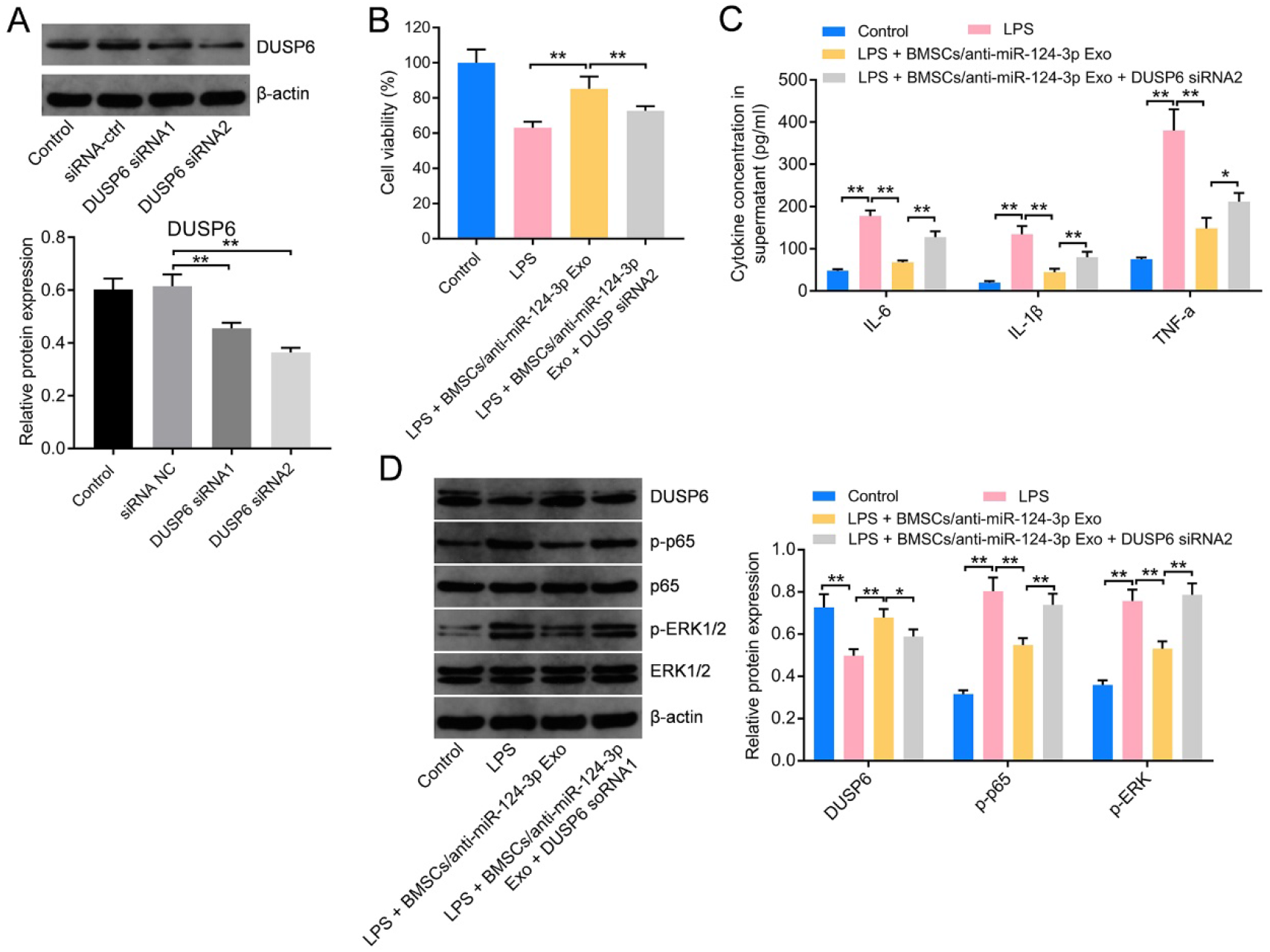
BMSCs/anti-miR-124-3p Exo alleviates LPS-induced inflammation of HEEC by upregulating DUSP6 and downregulating p-p65 and p-ERK. **(A)** The expression of DUSP6 in HEEC was assessed using western blot. **(B)** The viability of HEEC was assessed using CCK-8. **(C)** The expressions of IL-6, IL-1β and TNF-α in HEEC were assessed using ELISA. **(C)** The expressions of DUSP6, p-p65, p65, p-ERK1/2 and ERK1/2 in HEEC were assessed using ELISA.

### BMSCs/anti-miR-124-3p Exo protects against LPS-induced endometritis *in vivo* by upregulating DUSP6 and downregulating p-p65 and p-ERK

Finally, to investigate the roles of BMSCs/anti-miR-124-3p Exo on of endometritis procession *in vivo*, the mouse model of endometrial injury was constructed. The results of RT-qPCR indicated that LPS promoted the miR-124-3p expression in tissue, while this promotion was reserved by BMSCs/anti-miR-124-3p Exo (Fig. 6A). Additionally, HE and Masson staining showed that LPS induced endometritis, and this damage was relieved by BMSCs/anti-miR-124-3p Exo (Fig. 6B). Besides, LPS increased the expressions of IL-6, TNF-α and IL-1β in tissue, and those increases were reserved by BMSCs/anti-miR-124-3p Exo (Fig. 6C). Congruously, LPS downregulated the level of DUSP6 in tissue, and upregulated p-p65 and p-ERK levels; while the effects of LPS were reserved by BMSCs/anti-miR-124-3p Exo. Collectively, BMSCs/anti-miR-124-3p Exo protected against endometritis induced by LPS *in vivo* by upregulating DUSP6 and downregulating p-p65 and p-ERK.

**Figure 6.**
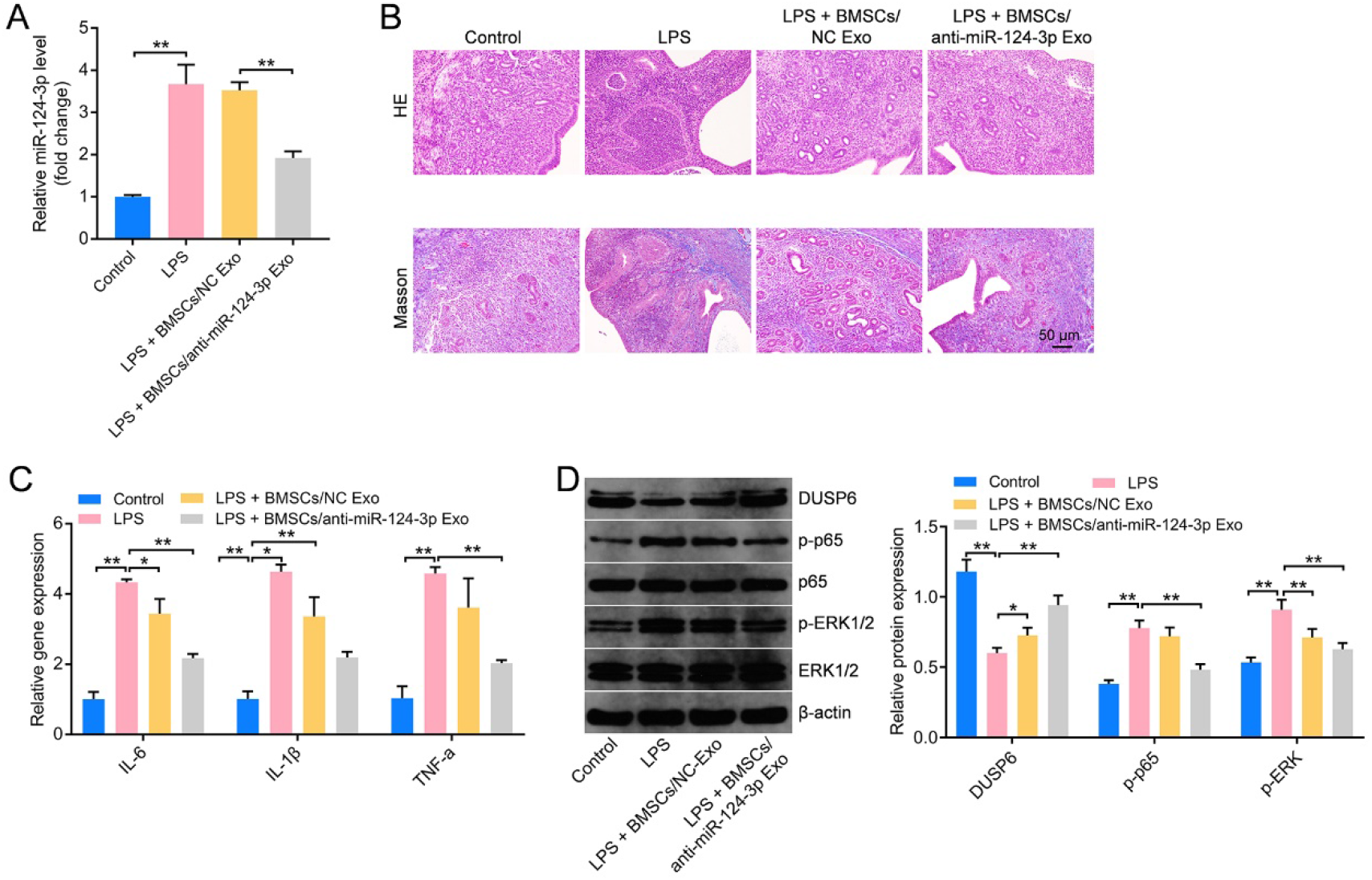
BMSCs/anti-miR-124-3p Exo protects against LPS-induced endometritis *in vivo* by upregulating DUSP6 and downregulating p-p65 and p-ERK. **(A)** The expression of miR-124-3p in tissue was assessed using RT-qPCR. **(B)** HE and Masson staining were performed to observe LPS induced endometritis in tissue. **(C)** The expressions of IL-6, IL-1β and TNF-α in tissue were assessed using ELISA. **(D)** The expressions of DUSP6, p-p65, p65, p-ERK1/2 and ERK1/2 in tissue were assessed using ELISA.

## Discussion

Endometritis is an inflammatory change in the endometrial structure caused by various reasons^1–3^. It has been reported that miRNAs can regulate cell growth and tissue differentiation. Therefore, miRNAs widely contributed to the development of organisms and the occurrence of diseases like endometritis^19,27,28^. For example, LPS promoted the expressions of IL-6, TNF-α and IL-1β in human HEECs, and the effect of LPS was inhibited by miR-643^29^. In addition, severe pathological changes occurred in LPS-induced endometritis mice, which were inhibited by the overexpression of miR-19a^30^. The overexpression of miR-19a could inhibit the expressions of IL-6, TNF-α, IL-1β and p-p65 in bEECs^30^. In the current study, LPS promoted the expressions of IL-6, TNF-α and IL-1β in HEEC, and those promotions were reversed through miR-124-3p inhibitor. Our results are consistent with these studies.

It has been reported that BMSCs can secrete a variety of cytokines and growth factors to enhance the differentiation and proliferation of hematopoietic stem cells^31^. Additionally, effects of anti-inflammatory, immunomodulatory and tissue repair are also belonged to MSCs^32^. For example, BMSC-separated exosome alleviates TGF-β1-induced the damage of the endometrium^24^. Besides, exosomes derived from human umbilical cord derived mesenchymal stem cells (UCMSCs) could deliver miR-7162-3p to endometrial stromal cell^33^. Exosomal miR-7162-3p could suppress the injury of endometrium that was induced by mifepristone ^33^. In the present study, exosomes could transfer miR-124-3p from BMSCs to HEEC. Additionally, BMSCs/anti-miR-124-3p Exo alleviated LPS-induced inflammation of HEEC.

## Conclusion

Our findings verified BMSCs/anti-miR-124-3p Exo protected against LPS-induced endometritis *in vitro* and *in vivo* by upregulating DUSP6 and downregulating p-p65 and p-ERK. Our findings add more value that BMSCs/anti-miR-124-3p Exo might be a promising new alternative for endometritis treatment.

## Acknowledgements

Not applicable.

## Author contributions

JW conceived the study and designed the experiments; YC and SZ completed the experiment and wrote the manuscript; XZ, YZ and SY discussed the results; JW and YC analyzed the data. YC, SZ and JW revised the manuscript.

## Availability of data and materials

The dataset used and/or analyzed in this study is available from the corresponding author on reasonable request.

